# Genetic Mapping Reveals a Major Locus for Boron Exclusion in Wild *Vitis*

**DOI:** 10.64898/2026.10.01.756076

**Authors:** Yaniv Lupo, Sadikshya Sharma, Jose Ramon Munoz, Veronica Nunez, Ana Gaspar, Noe Cochetel, Dario Cantu, Andrew J. McElrone, Luis Diaz-Garcia

## Abstract

Boron toxicity constrains grapevine (*Vitis*) production in irrigated arid and semi-arid regions, yet most available rootstocks are sensitive to excessive boron. Although underutilized *Vitis* species may harbor sources of adaptation, the genetic basis of natural variation in boron exclusion remains poorly understood, limiting the development of boron-tolerant rootstocks. We screened 267 wild *Vitis* accessions representing 16 species for their ability to limit leaf boron accumulation under excess boron conditions. Leaf boron concentration varied 190-fold among accessions and was highly heritable (entry-mean H² = 0.71). Its correlation with stomatal conductance was modest (r = 0.34), indicating that transpiration rate explained only a small proportion of the variation in leaf boron accumulation. Leaf boron was correlated with leaf chloride (r = 0.36), but their largely non-overlapping association profiles supported distinct genetic control. Genome-wide association analysis identified a suggestive association on chromosome 11. Independent quantitative trait locus mapping in a full-sib F_1_ population (n = 158) derived from Longii-9018 (*V. acerifolia*) × GRN3 (commercial rootstock) identified qBEx11.1 in the same syntenic region on chromosome 11, which explained 19.5% of the phenotypic variance. Positional candidate genes near the association peak included a pectinesterase and a MATE efflux transporter, suggesting roles for cell wall modification and membrane transport. Leaf spectral reflectance was associated with leaf boron concentration in both populations, particularly at visible and red-edge wavelengths, indicating potential for high-throughput phenotyping. Together, these findings identify qBEx11.1 as a priority locus for boron exclusion and support integrated genetic and spectral approaches for breeding boron-tolerant grapevine rootstocks.

**Plain Language Summary:** Boron is needed by plants in small amounts, but too much can damage them, especially in dry regions irrigated with boron-rich water. Grapevine rootstocks, the root systems onto which fruiting grape varieties are grafted, may help limit boron uptake. We screened 267 wild grapevines and offspring from two parent vines for their ability to limit boron buildup in leaves. Leaf boron levels differed nearly 190-fold. Genetic analyses of both groups identified the same region of chromosome 11 as important for limiting leaf boron. Transpiration rate explained only a small part of this variation, and boron and chloride buildup appeared to be controlled largely by different genetic factors. Measurements of how leaves reflected light also detected boron-related changes and may allow faster screening. These findings could help breeders develop grapevine rootstocks better suited to boron-affected soils and irrigation water.

## Introduction

Boron (B) is an essential plant micronutrient required for normal vegetative and reproductive development (Camacho-Cristóbal et al., 2008). Boron contributes to primary cell wall structure by forming borate diester cross-links between rhamnogalacturonan II (RG-II) molecules (Goldbach & Wimmer, 2007; O’Neill et al., 2004). Yet boron is unusual among mineral nutrients in that the concentration range separating deficiency from toxicity is exceptionally narrow (Brdar-Jokanović, 2020). This narrow range makes crops vulnerable to excess boron, particularly in irrigated arid and semi-arid regions, where naturally boron-rich soils and boron-containing irrigation water can result in sustained accumulation in plant tissues (Nable et al., 1997). Genetic improvement therefore offers a potentially durable strategy for retaining productivity in boron-affected environments.

In soil solution, boron occurs mainly as uncharged boric acid (H_3_BO_3_), which can enter roots by passive diffusion (Miwa & Fujiwara, 2010). In addition to this passive pathway, regulated root uptake, xylem loading, and distribution of boron within the plant involve nodulin 26-like intrinsic protein (NIP) channels and BOR-family efflux transporters (Takano et al., 2002, 2006; Yoshinari & Takano, 2017). Once in the xylem, boron moves to the shoot largely through the transpiration stream. Consequently, transpiration rate and stomatal conductance (gₛ) can influence boron delivery to the leaves, whereas boron toxicity can reduce g_s_, total transpiration, and root-to-shoot water transport (Macho-Rivero et al., 2018). Since boron has restricted phloem mobility, it tends to accumulate in older leaves and at leaf tips and margins (Brown & Shelp, 1997; Miwa & Fujiwara, 2010). Toxicity symptoms therefore often first appear at these sites as chlorosis and necrosis. Excess boron levels can also induce oxidative stress (Cervilla et al., 2007) and reduce growth and yield (Nable et al., 1997).

Grapevine (*Vitis*) rootstocks influence water and mineral uptake and root-to-shoot transport in grafted vines (Abbaspour et al., 2014; Dag et al., 2015; Fisarakis et al., 2001) and may help manage boron accumulation. Commercial rootstock breeding has drawn primarily on the North American species *V. riparia*, *V. rupestris*, and *V. berlandieri* (Heinitz et al., 2019). North American wild *Vitis* harbors extensive genetic and physiological diversity, including variation in water relations and gas exchange (Padgett-Johnson et al., 2003), adaptation to contrasting environments (Morales-Cruz et al., 2021), and exclusion of potentially toxic ions (Heinitz et al., 2020; Sharma et al., 2025). Previous studies of boron toxicity in grapevine have shown that increasing boron supply can raise leaf boron concentration, cause marginal chlorosis and necrosis, and reduce growth (Yermiyahu et al., 2007). Other reported responses include increased stomatal resistance (i.e., reduced gₛ), membrane damage, and oxidative stress (Gunes et al., 2006). Genotypes that maintain low leaf boron concentration under excess supply are described as boron excluders (Pech et al., 2013). A recent evaluation of 16 *Vitis* genotypes under excess boron found approximately 4.7-fold variation in leaf boron accumulation, displaying considerable natural variation in boron exclusion within the genus (Lupo et al., 2025). However, genotypic differences in growth under excess boron are not always explained by leaf boron concentration. Across 26 *Vitis* genotypes, shoot growth under excess boron ranged from 14% to 88% of the control but did not correlate with leaf boron (Pech et al., 2013).

The genetic architecture of boron exclusion remains poorly characterized in *Vitis*. In other species, genetic mapping has linked natural variation in boron accumulation to known transport genes. Genome-wide association mapping in barley (*Hordeum vulgare*) implicated the boric-acid channel *HvNIP2;2/HvLsi6* at a major-effect locus for grain boron concentration (Jia et al., 2021), while combined association and linkage mapping in Arabidopsis (*Arabidopsis thaliana*) identified variation in *BOR1* and *BOR2* associated with leaf boron accumulation (Tergemina et al., 2026). To our knowledge, no comparable genetic mapping study has been reported in grapevine. Phenotyping also presents a challenge because quantifying leaf boron requires destructive and labor-intensive elemental analysis, which tends to limit the number of samples and time points evaluated. Leaf spectral reflectance has demonstrated potential for nondestructive assessment of chloride (Cl) and boron in grapevine (Lupo et al., 2025; Sharma et al., 2024), but the consistency of boron-associated spectral signals across genetically distinct populations and their development during boron exposure remain unknown.

The relationship between boron and chloride (Cl^-^) is also relevant to rootstock breeding. Like boron, chloride is transported to the shoot through the xylem and can become toxic when it accumulates excessively in the leaves (Geilfus, 2018). Excess boron and salinity frequently co-occur in irrigated arid soils, and salinity can alter boron uptake and toxicity, although the direction of this effect varies among species and environmental conditions (Pandey et al., 2019; Yermiyahu et al., 2007). Wild *Vitis* contains substantial variation in chloride exclusion (Heinitz et al., 2020), and a major locus associated with chloride exclusion was identified using an overlapping diversity panel and validated in a biparental family (Sharma et al., 2025). Whether genotypes selected for low chloride accumulation also accumulate less boron therefore remains an important practical question for rootstock breeding.

To address these gaps, we characterized boron exclusion under excess boron by measuring leaf boron concentration, g_s_, and photosystem II quantum yield (Φ_PSII_) across a wild *Vitis* diversity panel and a full-sib F_1_ population. We combined genome-wide association analysis in the diversity panel with QTL mapping in the F_1_ population to identify genomic regions associated with variation in leaf boron accumulation under boron stress. We further compared leaf boron and chloride concentrations within the diversity panel and evaluated leaf spectral reflectance as a high-throughput phenotyping approach in both populations. We hypothesized that leaf boron exclusion would be heritable and associated with detectable genomic loci in wild *Vitis*, and that variation in gₛ would explain only part of the genotypic variation in leaf boron accumulation.

## Materials and Methods

### Plant material and experimental design

Two grapevine populations were used in this study: a wild *Vitis* diversity panel of 267 accessions representing 16 species, and a full-sib F_1_ population of 162 progeny derived from a cross between Longii-9018 (*V. acerifolia* wild selection) and GRN3 (commercial rootstock; *V. rufotomentosa* × (Dog Ridge × Riparia Gloire) × *V. champinii*). Most accessions in the diversity panel originated from the southwestern United States and are maintained in the University of California, Davis grapevine germplasm collection. The following species were included: *V. acerifolia*, *V. aestivalis*, *V. arizonica*, *V. berlandieri*, *V. californica*, *V. candicans*, *V. champinii*, *V. × doaniana*, *V. girdiana*, *V. labrusca*, *V. monticola*, *V. riparia*, *V. rupestris*, *V. treleasei*, *V. cinerea*, and *V. vulpina*. The diversity panel was evaluated in two independent trials, with up to four replicates per accession (a mean of 3.9 total replicates per accession). The F_1_ population was evaluated in one trial with four replicates per progeny. Sample sizes for each analysis are summarized in Table S1. Plants were propagated from green cuttings and grown in 1-L pots in a pad-and-fan-cooled greenhouse at the University of California, Davis. All plants were fertigated daily with a complete nutrient solution described by Lupo et al. (2025). The diversity panel was fertigated with the base nutrient solution adjusted to a final boron concentration of 12 mg B L⁻^1^ (supplied as H_3_BO_3_) for 27 days in trial 1 and 28 days in trial 2. The F_1_ population was fertigated with the same base nutrient solution adjusted to a final boron concentration of 20 mg B L⁻^1^ (supplied as H_3_BO_3_) for 22 days. Both parents of the F_1_ population had shown relatively low leaf boron accumulation in a previous screening experiment under excess boron (Lupo et al., 2025); therefore, a higher boron concentration was used to increase phenotypic differentiation among their progeny. Representative boron-toxicity symptoms observed in the F_1_ population at harvest are shown in Figure S1.

### Leaf boron and physiological measurements

Leaves were destructively sampled at the end of each experiment. All leaves from each plant were combined, oven-dried to constant weight, and ground to a fine powder. Three replicates from each accession/progeny were randomly selected for boron analysis. Leaf boron concentrations were determined at the UC Davis Analytical Laboratory using inductively coupled plasma atomic emission spectrometry (ICP-AES) following microwave digestion of 400 mg of dry tissue with nitric acid and hydrogen peroxide. Leaf boron concentrations were expressed on a dry-weight basis (mg B kg⁻^1^ DW).

Stomatal conductance and Φ_PSII_ were measured on fully expanded, sun-exposed leaves between 1100 and 1300 h using a LI-600 porometer/fluorometer (LI-COR Biosciences, Lincoln, NE). Raw negative g_s_ values were excluded before model fitting as instrument artifacts, and slightly negative g_s_ BLUEs arising from model adjustment were retained.

### Phenotypic data analysis

Statistical analyses were performed in R v4.4.1 (R Core Team, 2024). Best linear unbiased estimates (BLUEs) were generated separately for each trait using the following model:

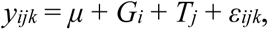

where *yᵢⱼ_k_* represents the phenotype of replicate *k* of genotype *i* in trial *j*; *μ* is the overall mean; *Gᵢ* is the fixed effect of genotype *i*; *Tⱼ* is the fixed effect of the independent trial or laboratory batch in the diversity panel and F_1_ population, respectively; and *εᵢⱼ_k_* is the residual error. BLUEs were extracted with the emmeans package v2.0.1 (Lenth and Piaskowski, 2025). Species represented by at least three accessions with non-missing data were included in species-level comparisons, and only species meeting this criterion for both g_s_ and Φ_PSII_ were included in the physiological-trait comparisons (Figure 1D-F). Species differences were tested using Kruskal-Wallis tests followed by Dunn’s pairwise comparisons with Benjamini-Hochberg (BH)-adjusted two-sided P values (α = 0.05; Benjamini & Hochberg, 1995).

**Figure 1.**
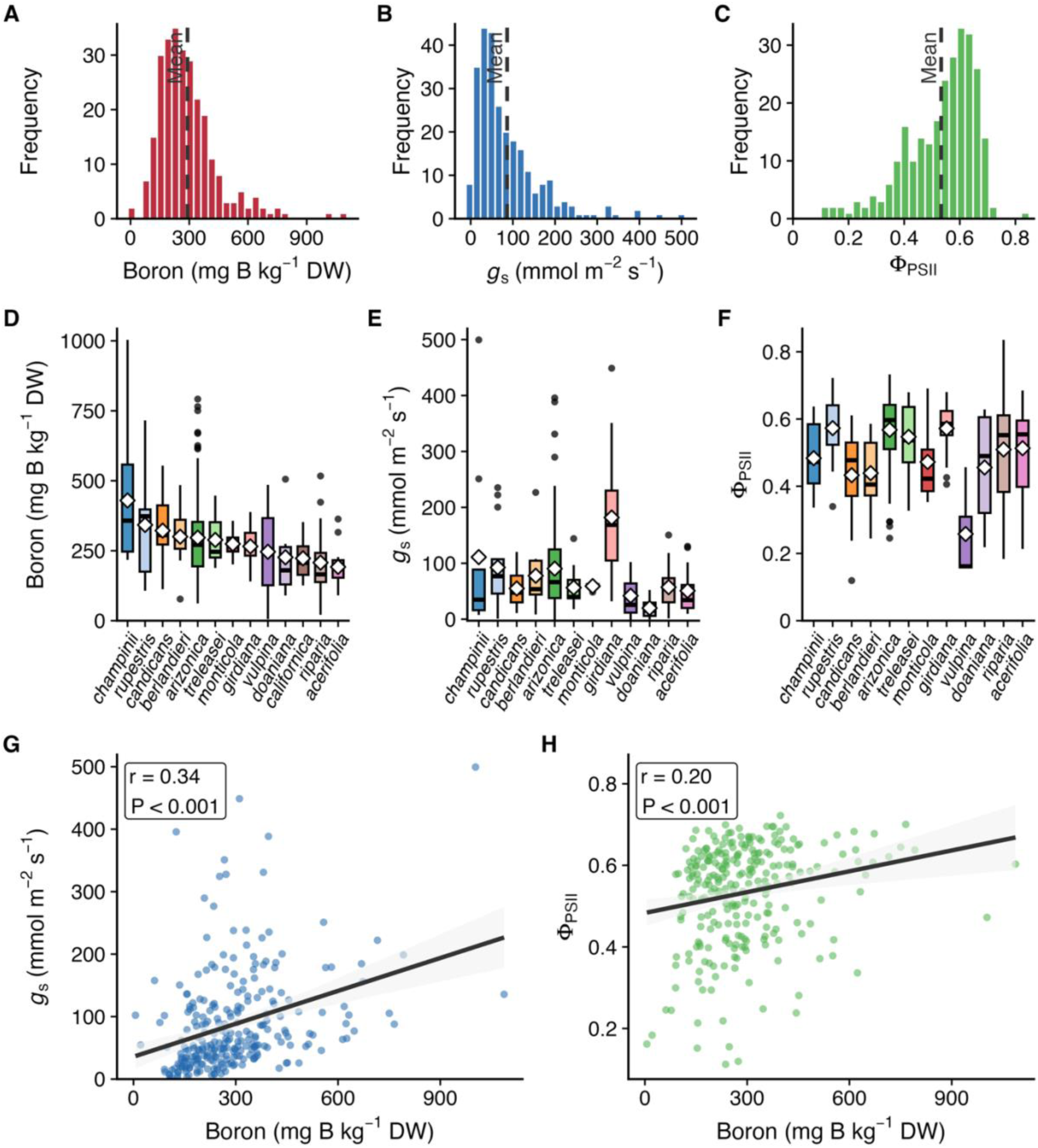
Phenotypic variation in leaf boron accumulation and physiological traits across the wild *Vitis* diversity panel. Frequency distributions of BLUEs for (A) leaf boron concentration (n = 267), (B) stomatal conductance (gₛ; n = 265), and (C) photosystem II quantum yield (Φ_PSII_; n = 265). Dashed vertical lines indicate the trait means. Species-level variation for (D) leaf boron concentration, (E) gₛ, and (F) Φ_PSII_. White diamonds indicate species means. Only species represented by at least three accessions with non-missing data were included in each species-level panel. (G) Pearson correlations between leaf boron and gₛ (n = 265), and (H) leaf boron and Φ_PSII_ (n = 265). Regression lines and 95% confidence intervals are shown.

For heritability estimation, linear mixed models were fitted by restricted maximum likelihood (REML) using the lme4 package v1.1.38 (Bates et al., 2015). Genotype was treated as a random effect in both populations, with trial included as a fixed effect for the diversity panel and laboratory batch as a fixed effect for the F_1_ population. Broad-sense heritability on an individual-plant basis was calculated as:

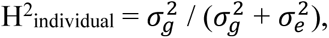

and broad-sense heritability on an entry-mean basis was calculated as:

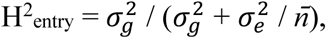

where 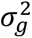 and 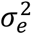 are the genotypic and residual variance components, respectively, and *n̄* is the mean number of observations per accession or progeny (Holland et al., 2003).

### Genome-wide association analyses

Sequencing data were compiled from previously published studies (Cochetel et al., 2023; Morales-Cruz et al., 2021, 2023) and are publicly available in the NCBI Sequence Read Archive under BioProject IDs PRJNA731597, PRJNA842753, and PRJNA984685. Raw reads were filtered using Trimmomatic v0.36 (Bolger et al., 2014) and quality-checked with FastQC v0.11.9 (Andrews, 2010). Reads were scanned using a four-base sliding window and trimmed when the average base quality fell below Q20; leading and trailing bases below Q3 were removed and only reads ≥ 60 bp were retained. Filtered reads were aligned to haplotype 1 of the *Vitis girdiana* SC2 genome v1.0 (Cochetel et al., 2023) using BWA-MEM (BWA v0.7.12-r1039; Li, 2013).

Joint genotyping was performed using GATK v4.0.12.0 (Van der Auwera & O’Connor, 2020), following the pipeline described in Cochetel et al. (2023). Briefly, duplicate reads were marked using Picard MarkDuplicates (v2.8.2-4-g2105a1e-SNAPSHOT), and read groups were added with AddOrReplaceReadGroups. Variant calling was performed with GATK HaplotypeCaller using a ploidy of 2 and a minimum base quality of Q20, followed by joint genotyping with GenotypeGVCFs. SNP filtering was conducted with bcftools v1.19 (Danecek et al., 2021) using GATK hard-filtering criteria (QD < 2 || FS > 60 || SOR > 3 || MQ < 40 || MQRankSum < −12.5 || ReadPosRankSum < −8). Only biallelic SNPs with <25% missing data and a minor allele frequency (MAF) > 0.10 were retained for association analyses.

Genome-wide association analyses were performed in GEMMA v0.98.3 (Zhou & Stephens, 2012) using a mixed linear model. A standardized genomic relationship matrix was calculated in GEMMA using -gk 2. The first five principal components, calculated in PLINK v1.9 (Chang et al., 2015) from the same marker set used for GWAS, were included as fixed-effect covariates to account for population structure. For the primary leaf boron analysis, g_s_ BLUEs were additionally included as a covariate to account for variation in leaf boron associated with transpiration. To assess whether the chromosome 11 association depended on this adjustment, the leaf boron GWAS was repeated using the same accessions, SNPs, genomic relationship matrix, and five principal components but without gₛ as a covariate.

Bonferroni-corrected genome-wide significance thresholds (P < 2.81 × 10⁻⁸ for leaf boron and P < 2.84 × 10⁻⁸ for g_s_) and a suggestive threshold of P < 1 × 10⁻⁵ were used to identify marker-trait associations. Lead SNPs were defined as the most significant SNP within non-overlapping 500-kb genomic windows. All genomic coordinates are reported relative to haplotype 1 of the *V. girdiana* SC2 reference genome.

Candidate genes were identified using the VITVgdSC2_v1.0 annotation for haplotype 1 of the *V. girdiana* SC2 reference genome, available through grapegenomics.com. Candidate-gene annotation focused on the chromosome 11 boron lead SNP and three selected gₛ lead SNPs shown in Figure 2H-J (chr18:5.94 Mb, chr3:2.55 Mb, and chr11:2.14 Mb). Genes located within ±50 kb of these four lead SNPs were identified, and genes with predicted functions potentially related to boron transport, ion transport, boron accumulation, or stomatal regulation were highlighted. For each gene, we compiled the gene ID, genomic coordinates, strand, signed distance and orientation relative to the lead SNP, whether the lead SNP was located within the gene, and the corresponding functional annotation. All genes within these four intervals are reported in Table S2.

**Figure 2.**
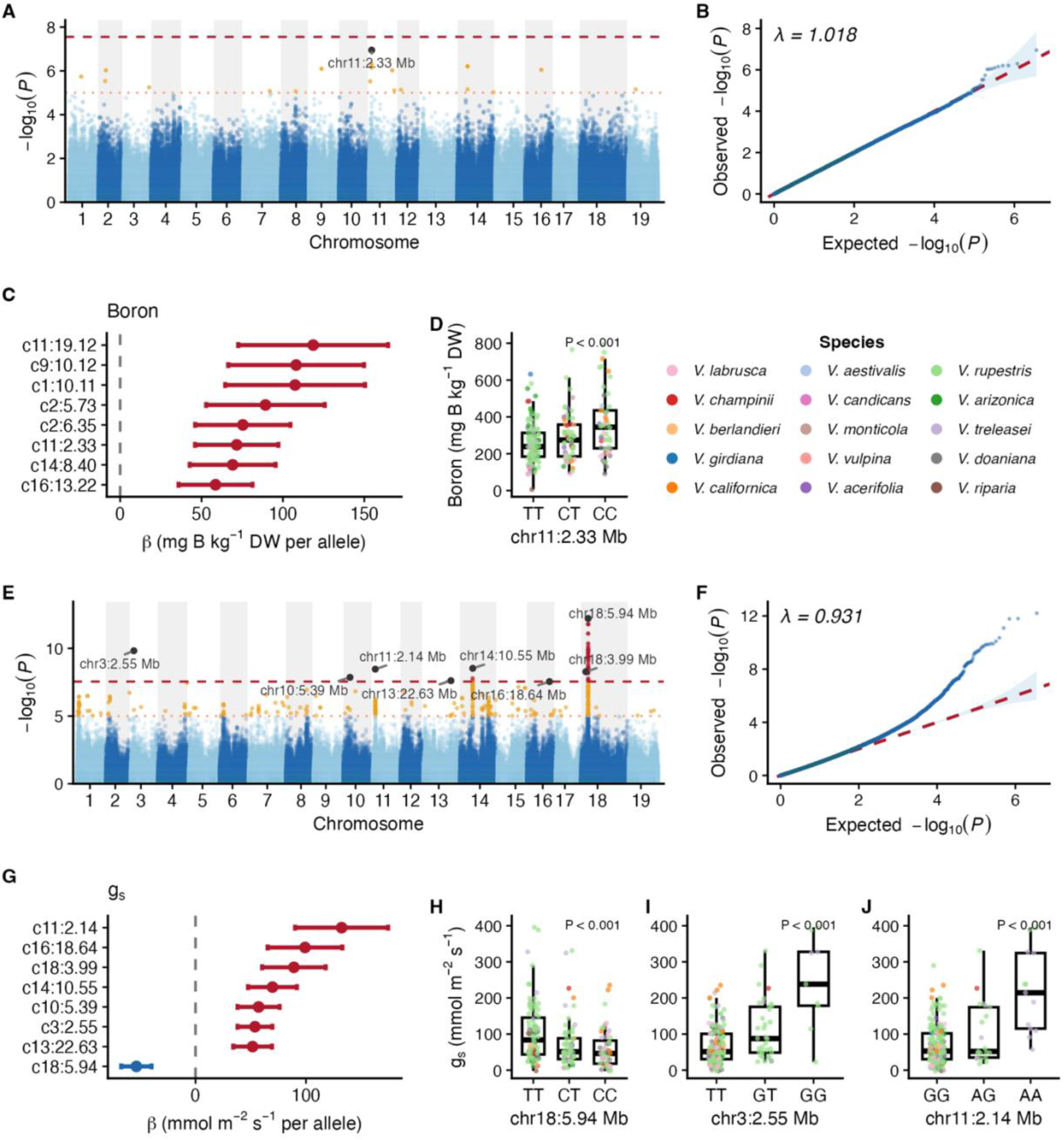
Genome-wide association analyses of leaf boron concentration and stomatal conductance (gₛ) in the wild *Vitis* diversity panel. (A) Manhattan plot for leaf boron concentration. (B) Quantile-quantile plot for the leaf boron GWAS. (C) Effect estimates for the eight lead suggestive boron-associated loci. (D) Leaf boron BLUEs by genotype class at chr11:2.33 Mb. (E) Manhattan plot for gₛ. (F) Quantile-quantile plot for the gₛ GWAS. (G) Effect estimates for the eight lead Bonferroni-significant gₛ-associated loci. (H-J) gₛ BLUEs by genotype class at three Bonferroni-significant loci: (H) chr18:5.94 Mb, (I) chr3:2.55 Mb, and (J) chr11:2.14 Mb. In A and E, the dotted and dashed horizontal lines indicate the suggestive threshold (P < 1 × 10⁻⁵) and the trait-specific Bonferroni thresholds (P < 2.81 × 10⁻⁸ for leaf boron and P < 2.84 × 10⁻⁸ for gₛ), respectively. In B and F, the diagonal line represents the expected null distribution. In C and G, points represent SNP effect estimates (β), and horizontal bars indicate 95% confidence intervals.

### QTL mapping

QTL mapping was performed in the Longii-9018 × GRN3 F_1_ population (n = 158 progeny with marker data) using leaf boron BLUEs. Genotyping-by-sequencing, variant calling, and linkage-map construction for this population were carried out as in Sharma et al. (2025), using the *V. acerifolia* Longii-9018 genome as the reference (Cochetel et al., 2023). Linkage maps were constructed using a pseudo-testcross mapping strategy (Grattapaglia & Sederoff, 1994). The Longii-9018 parental map, comprising markers segregating as AB × AA (lm × ll), contained 2,611 markers distributed across 740 unique genetic positions (bins), whereas the GRN3 parental map contained 4,842 markers across 784 bins. The merged genetic map comprised 1,038 markers (262 hk × hk, 426 lm × ll, and 350 nn × np; Figure S2A) and was analyzed as a four-way cross in the R/qtl package v1.74 (Broman et al., 2003). Genotype probabilities were calculated at 1-cM intervals using the Kosambi map function and a genotyping error rate of 0.001. QTL scans were performed using Haley-Knott regression (Haley & Knott, 1992). The genome-wide 5% logarithm of odds (LOD) threshold was determined from 1000 permutations, and the 1.5-LOD support interval was calculated in R/qtl. To evaluate each parental contribution separately, parent-specific QTL scans were performed for Longii-9018 and GRN3, with each parental map analyzed as a backcross (Figure S2B, C).

The percentage of phenotypic variance explained by the QTL was calculated as 100 × R^2^, where R^2^ was obtained from a one-way ANOVA comparing leaf boron BLUEs among genotype classes at the marker corresponding to the maximum LOD score. Pairwise differences among genotype classes were evaluated using Tukey’s HSD. At this marker, the effects of the Longii-9018 haplotype (A versus B), the GRN3 haplotype (C versus D), and their interaction were tested using two-way ANOVA.

Because the molecular marker sets for the two populations were generated using different approaches and different reference genomes for variant calling (diversity panel: whole-genome sequencing aligned to the *V. girdiana* SC2 reference genome; Longii-9018 × GRN3 F_1_ population: genotyping-by-sequencing aligned to the Longii-9018 reference genome), haplotype 1 assemblies of the two references were aligned using minimap2 v2.31 (Li, 2018). This alignment was then used to translate genomic coordinates between reference assemblies and enable direct comparison of the GWAS and QTL signals identified in the two populations.

### Boron-chloride phenotypic and GWAS comparison

The diversity panel was previously phenotyped for leaf chloride concentration by Sharma et al. (2025). Chloride was measured in dried leaf tissue using a silver-ion titration chloridometer (Model 926, Nelson-Jameson Inc.). Leaf boron and chloride concentrations were correlated among 241 accessions with data for both traits using Pearson correlation. For the genetic comparison, both GWAS were rerun in GEMMA using the same 222 accessions, 1,703,430 SNPs, genomic relationship matrix, and five principal components. Trait-specific BLUEs were re-estimated, and gₛ was excluded from the matched boron GWAS to maintain the same model structure for both traits. Summary statistics were merged by chromosome and position. For SNPs reaching P < 1 × 10⁻⁵ for either trait, chloride effect estimates were aligned to the boron GWAS effect allele before effect directions were compared. To summarize the genome-wide distribution of associations, suggestive SNPs were grouped into non-overlapping 500-kb windows, and the most significant SNP for each trait in each window was retained.

### Leaf spectral reflectance analysis

Leaf spectral reflectance was measured using a CI-710s leaf spectrometer (CID Bio-Science, Camas, WA; 2.4 nm spectral resolution) on the fourth leaf from the base of each plant. Measurements were collected between 0900 and 1200 h for the diversity panel and between 1200 and 1500 h for the F_1_ population. Each leaf was measured once, and white and dark reference calibrations were performed at the start of each measurement day. Quality control and preprocessing were carried out in the waves v0.2.6 package (Hershberger et al., 2021). Spectra were interpolated to 1-nm resolution over 380-1,100 nm, yielding 721 wavelengths. Scans with reflectance at 1,090 nm outside 20-75%, or with scores more than three standard deviations from the mean on either of the first two principal-component axes, were excluded. Raw reflectance spectra were retained for visualization, whereas Pearson correlations between genotype-mean SNV-transformed reflectance and leaf boron BLUEs were calculated at each wavelength. P values were adjusted separately for each population and time point using the BH-FDR and Bonferroni procedures. Low- and high-boron groups were defined as the lower and upper 20% of retained genotypes based on leaf boron BLUEs. High-low spectral differences were calculated from raw genotype-mean reflectance as the mean reflectance of the high-boron group minus that of the low-boron group.

In the diversity panel, reflectance was measured once per plant at 26 and 21 days after treatment initiation in trials 1 and 2, respectively. Raw spectra were averaged within accession and trial and then across trials. For correlation analysis, spectra were mean-centered by wavelength within each trial, SNV-transformed, and averaged using the same procedure. In the F_1_ population, reflectance was measured before treatment and at 5, 13, and 21 days after treatment initiation (day of treatment; DOT). Quality control and SNV transformation were performed separately at each time point, and spectra were then averaged by progeny. The DOT 13 measurement was used for comparisons with the diversity panel.

## Results

### Leaf boron accumulation and physiological traits vary widely across the wild *Vitis* diversity panel

Across the 267-accession diversity panel, representing 16 *Vitis* species, substantial within- and among-species variation was observed for leaf boron concentration, g_s_, and Φ_PSII_. Leaf boron BLUEs ranged from 5.7 to 1,087.2 mg B kg⁻¹ DW (approximately 190-fold), with a mean of 291 mg B kg⁻¹ DW (Figure 1A). BLUEs for g_s_ ranged from −2.6 to 499.4 mmol m⁻² s⁻¹ (mean = 86.4 mmol m⁻² s⁻¹; Figure 1B). BLUEs for Φ_PSII_ ranged from 0.11 to 0.84 (mean = 0.53; Figure 1C).

Leaf boron concentration differed significantly among species (Kruskal-Wallis H = 27.92, P = 5.68 × 10⁻^3^). *V. arizonica*, *V. candicans*, and *V. champinii* had significantly higher values than *V. acerifolia* and *V. riparia* (BH-adjusted Dunn tests, P < 0.05; Figure 1D). Stomatal conductance also differed among species (H = 39.06, P = 5.17 × 10⁻^5^); *V. girdiana* had significantly higher g_s_ than *V. acerifolia*, *V. arizonica*, *V. candicans*, *V. champinii*, *V. doaniana*, *V. riparia*, and *V. treleasei*, while *V. arizonica* had higher values than *V. doaniana* (BH-adjusted Dunn tests, P < 0.05; Figure 1E). Φ_PSII_ differed among species (H = 40.50, P = 2.94 × 10⁻^5^); *V. arizonica* had significantly higher values than *V. berlandieri* and *V. candicans*, while *V. girdiana* had higher values than *V. candicans* (BH-adjusted Dunn tests, P < 0.05; Figure 1F). Leaf boron concentration was positively correlated with g_s_ (Pearson’s r = 0.34, P = 1.31 × 10⁻^8^; Figure 1G) and had a weaker correlation with Φ_PSII_ (r = 0.20, P = 8.16 × 10⁻^4^; Figure 1H). Entry-mean broad-sense heritability was 0.71 for leaf boron concentration and 0.74 for g_s_, with corresponding individual-plant estimates of 0.46 and 0.44, respectively.

### GWAS identifies a suggestive association for boron exclusion on chromosome 11

The leaf boron GWAS included 1,776,991 SNPs across 224 accessions. No SNP reached genome-wide significance at a Bonferroni genome-wide significance threshold of P < 2.81 × 10⁻⁸ (Figure 2A, B). Only 20 SNPs passed a suggestive threshold of P < 1 × 10⁻⁵; grouping these SNPs into non-overlapping 500-kb windows identified eight lead suggestive loci, with estimated effects ranging from 58.6 to 118.6 mg B kg⁻¹ DW per copy of the minor allele (Figure 2C). The strongest association was located at chr11:2.33 Mb (P = 1.12 × 10⁻⁷; β = 71.6 mg B kg⁻¹ DW per copy of the minor allele; Figure 2D). The chromosome 11 association occurred at the same position in an analysis without gₛ as a covariate (P = 3.23 × 10⁻⁷; data not shown). Within the ±50-kb interval surrounding the chromosome 11 lead SNP, the two closest annotated genes encoded a COMPASS-like WDR5A component and a probable glutamate carboxypeptidase, located approximately 0.5 and 1.3 kb from the lead SNP, respectively. A probable pectinesterase 53 and a MATE efflux family protein were located approximately 13 and 33 kb from the lead SNP, respectively. All genes within the ±50-kb interval surrounding the chromosome 11 boron lead SNP are listed in Table S2.

### GWAS identifies eight loci associated with stomatal conductance

GWAS on gₛ included 1,758,640 SNPs across 233 accessions, of which 44 SNPs passed the Bonferroni genome-wide significance threshold of P < 2.84 × 10⁻⁸ (Figure 2E, F). Grouping these SNPs into non-overlapping 500-kb windows identified eight lead loci (Figure 2G). The strongest association was located at chr18:5.94 Mb (P = 6.19 × 10⁻¹³; β = −53.5 mmol m⁻² s⁻¹ per copy of the minor allele; Figure 2H). At chr3:2.55 Mb and chr11:2.14 Mb, the minor alleles were associated with higher gₛ (β = 53.8 and 132.2 mmol m⁻² s⁻¹, respectively), and genotype-class differences were significant (Figure 2I, J). The gₛ-associated SNP at chr11:2.14 Mb was located approximately 190 kb from the strongest boron-associated SNP at chr11:2.33 Mb. At chr18:5.94 Mb, the lead SNP occurred within a gene encoding a GATA transcription factor, and a second GATA-type zinc-finger gene was located approximately 17 kb away. At chr3:2.55 Mb, the lead SNP was located approximately 1.6 kb from an L-ascorbate oxidase homolog. Genes located within ±50 kb of the three selected gₛ lead SNPs at chr18:5.94 Mb, chr3:2.55 Mb, and chr11:2.14 Mb are listed in Table S2.

### QTL mapping identifies a major chromosome 11 locus for boron exclusion

Leaf boron in the F_1_ population was continuously distributed among the 162 progeny and showed transgressive segregation; 46 progeny had lower values than GRN3, whereas 13 had higher values than Longii-9018 (Figure 3A). Entry-mean broad-sense heritability was 0.62, with an individual-plant estimate of 0.36.

**Figure 3.**
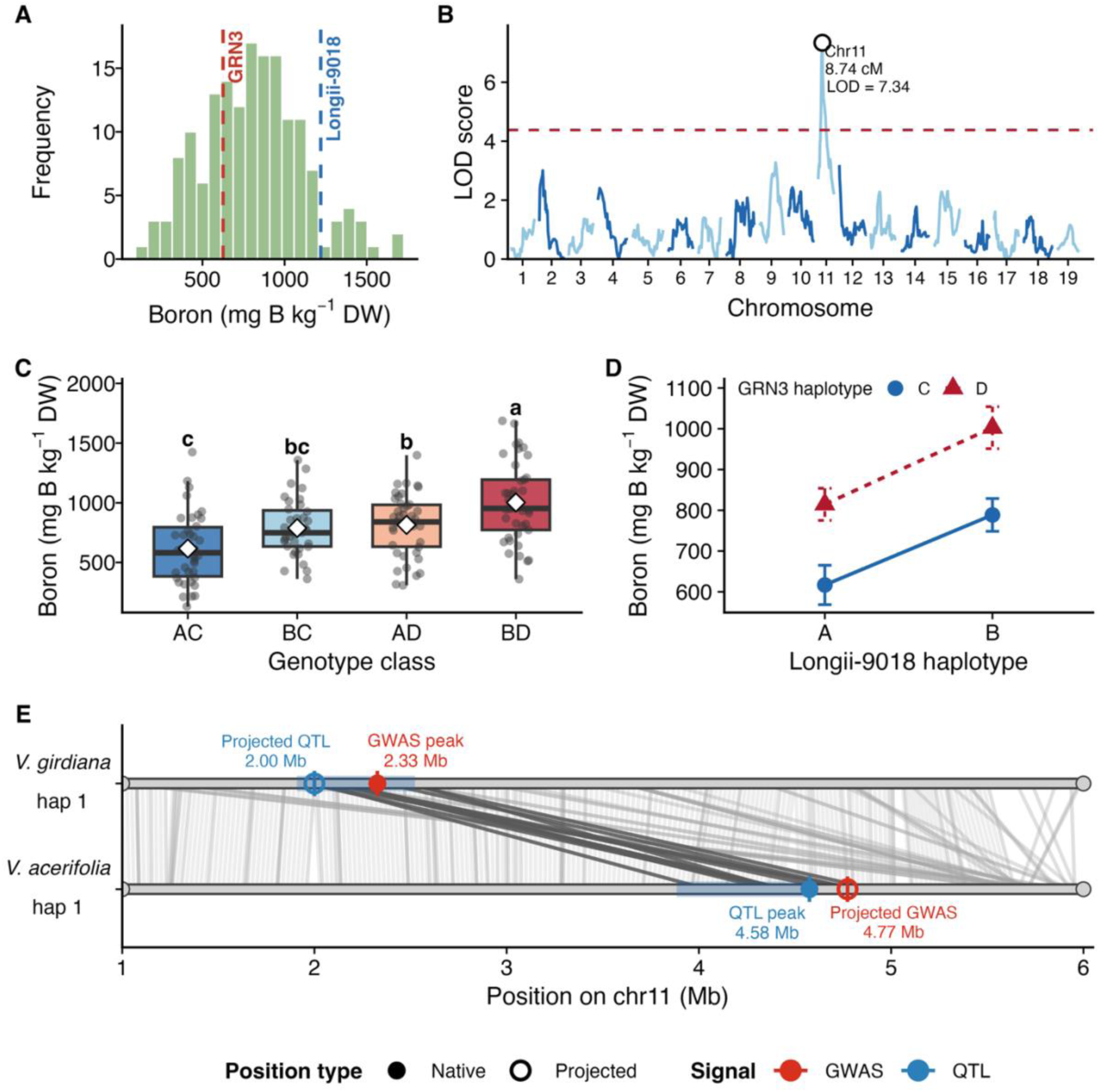
QTL mapping of leaf boron concentration in the Longii-9018 × GRN3 F_1_ population. (A) Frequency distribution of leaf boron BLUEs among 162 F_1_ progeny. Dashed vertical lines indicate the leaf boron BLUEs of the GRN3 and Longii-9018 parents. (B) Genome-wide LOD profile for leaf boron BLUEs. The dashed horizontal line indicates the 5% genome-wide LOD threshold (4.37). (C) Distribution of leaf boron BLUEs by genotype classes at the marker corresponding to the highest LOD score. White diamonds indicate genotype-class means. Genotype classes not sharing a letter differed significantly (Tukey-adjusted P < 0.05). (D) Haplotype means at the marker with the highest LOD score. Points represent mean leaf boron concentration, and error bars indicate ± SE. (E) Local sequence alignment of the chromosome 11 region containing the QTL and GWAS peaks between the *V. acerifolia* Longii-9018 (AC9018) and *V. girdiana* SC2 assemblies. Light gray lines represent broad alignment blocks across the displayed region, whereas dark gray lines represent the targeted local alignment used to project the peak positions. Filled and open symbols indicate native and projected positions, respectively. Blue bars indicate the portion of the 6.0-10.2-cM 1.5-LOD support interval represented within the displayed region of the Longii-9018 assembly and its projection onto SC2.

QTL mapping of leaf boron BLUEs from 158 progeny with marker data identified a significant QTL on chromosome 11, explaining 19.5% of the phenotypic variance (Figure 3B). The maximum LOD score of 7.34 occurred at an observed marker positioned at 8.74 cM, corresponding to 4.58 Mb on chromosome 11 of the *V. acerifolia* Longii-9018 assembly. This score exceeded the 5% genome-wide permutation threshold of 4.37, and the 1.5-LOD support interval spanned 6.0-10.2 cM (Figure S2D). Parent-specific scans detected significant chromosome 11 signals for both Longii-9018 and GRN3, indicating that allelic segregation from both parents was associated with leaf boron concentration, although their maximum LOD scores occurred at different positions (Figure S2B, C).

At the marker corresponding to the highest LOD score, genotype-class means were lowest for AC (616.8 mg B kg⁻¹ DW), highest for BD (1,002.7 mg B kg⁻¹ DW), and intermediate for BC and AD (788.6 and 814.6 mg B kg⁻¹ DW, respectively; Figure 3C). Mean leaf boron concentration was 179.9 mg B kg⁻¹ DW higher among progeny carrying the Longii-9018 B haplotype than among those carrying the A haplotype. The corresponding difference between the GRN3 D and C haplotypes was 205.9 mg B kg⁻¹ DW (Figure 3D). The parental haplotype effects were additive, with no evidence of interaction (P = 0.86; Figure 3D).

Alignment of chromosome 11 regions between the *V. girdiana* SC2 and *V. acerifolia* Longii-9018 assemblies placed the QTL and GWAS peaks within the same syntenic region (Figure 3E). The peaks were estimated to be approximately 330 kb apart on the SC2 assembly and 200 kb apart on the Longii-9018 assembly.

### Leaf boron and chloride concentrations are phenotypically correlated but show largely trait-specific GWAS signals

Leaf boron and chloride concentrations were positively correlated across the 241 accessions with measurements for both traits (Pearson’s r = 0.36, P = 1.03 × 10⁻⁸; Figure 4A). GWAS were then compared using the same 222 accessions and 1,703,430 SNPs. Of the SNPs reaching the suggestive threshold, 45 were associated only with boron and 73 only with chloride; none reached the threshold for both traits (Figure 4B). When effects were compared for the same allele in both GWAS, 116 of the 118 suggestive SNPs (98.3%) had estimated effects in the same direction for both traits, whereas two had opposing effects (Figure 4C). After retaining the most significant SNP within each 500-kb window, the boron associations represented 25 windows across 14 chromosomes, and the chloride associations represented 36 windows across 16 chromosomes. No 500-kb window contained a suggestive association for both traits (Figure 4D). Thus, despite the positive phenotypic correlation, the strongest GWAS signals were largely trait specific.

**Figure 4.**
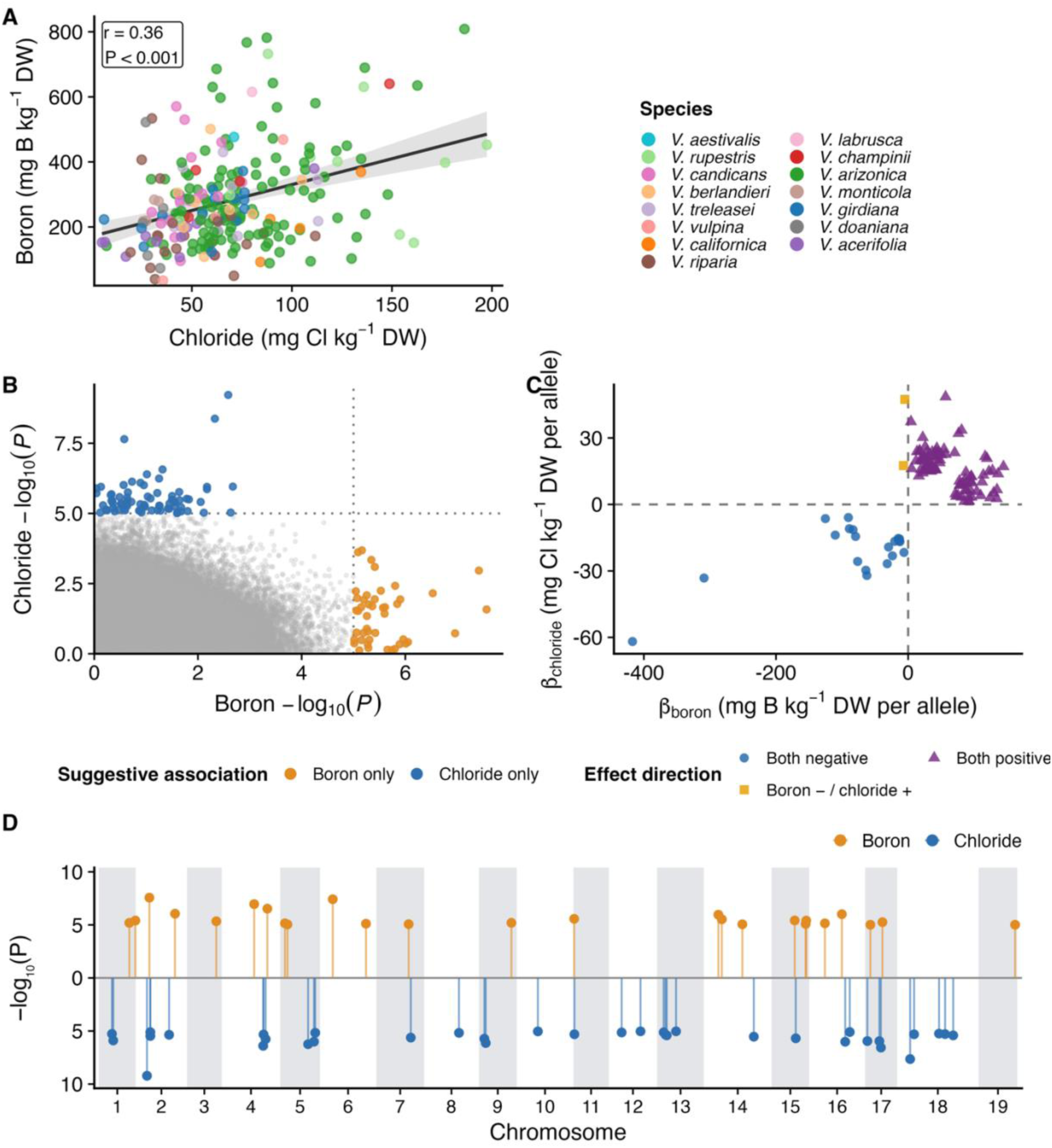
Phenotypic and genome-wide association comparisons of leaf boron and chloride concentrations in the wild *Vitis* diversity panel. (A) Pearson correlation between leaf chloride and boron concentrations (n = 241). Regression line and 95% confidence intervals are shown. (B) Comparison of association significance between GWAS of leaf boron and chloride concentrations (n = 222). Colored points represent SNPs associated only with boron or chloride. Dotted lines indicate the suggestive threshold of P < 1 × 10⁻⁵. (C) Comparison of estimated allelic effects for SNPs reaching the suggestive threshold for at least one trait (n = 118). (D) Mirrored genome-wide distribution of suggestive associations after retaining the most significant SNP within each non-overlapping 500-kb window. Boron associations are plotted above zero and chloride associations below zero for visualization. Points show −log₁₀(P) for the lead SNP in each window, and alternating gray shading distinguishes adjacent chromosomes.

### Leaf spectral reflectance correlates with leaf boron concentration in both populations

The diversity panel and F_1_ population showed similar mean reflectance profiles, characterized by low reflectance in the visible region, a sharp red-edge transition near 700 nm, and high reflectance in the near-infrared region (Figure 5A). Reflectance at numerous wavelengths was correlated with leaf boron concentration in both populations (Figure 5B). In the diversity panel, 244 of 721 wavelength-wise correlations remained significant after BH-adjustment. The strongest positive correlation occurred at 681 nm (r = 0.31), whereas the strongest negative correlation occurred at 719 nm (r = −0.28; Figure 5B, C). Significant wavelengths occurred primarily within 400-505, 530-575, 650-691, and 703-739 nm. In the F_1_ population, 520 of 721 wavelength-wise correlations remained significant after adjustment. The strongest positive and negative correlations occurred at 678 nm (r = 0.62) and 563 nm (r = −0.62), respectively. Significant wavelengths occurred primarily within 380-509, 514-644, 651-742, and 845-1,022 nm. Mean raw reflectance was compared between accessions or progeny in the lowest and highest 20% based on leaf boron concentration within each population. Low- and high-boron groups corresponded to BLUEs of ≤169.0 and ≥374.4 mg B kg⁻¹ DW in the diversity panel and ≤553.6 and ≥1,049.0 mg B kg⁻¹ DW in the F_1_ population. The two groups showed the largest reflectance differences across the visible and red-edge regions (Figure 5D).

**Figure 5.**
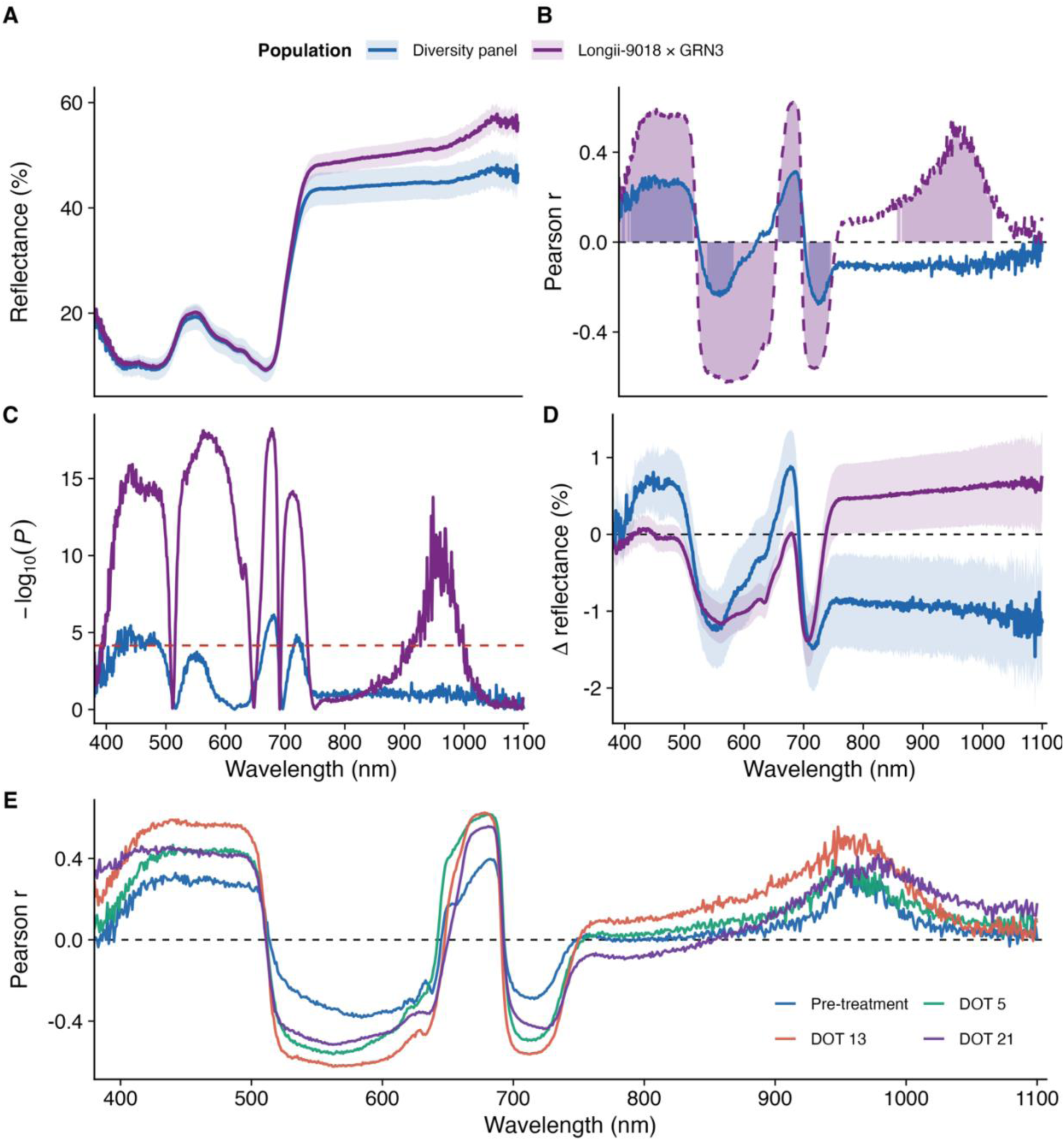
Associations between leaf spectral reflectance and leaf boron concentration in the wild *Vitis* diversity panel and the Longii-9018 × GRN3 F_1_ population. (A) Mean leaf reflectance spectra across 380-1,100 nm. Lines show population means, and shaded bands indicate ±1 SD (n = 238 and 162 for the diversity panel and the F_1_ population, respectively). (B) Wavelength-wise Pearson correlations between leaf reflectance and leaf boron concentration. Shaded regions identify wavelengths significant at a 5% false discovery rate (FDR). (C) Wavelength-wise significance profiles. The dotted red horizontal line indicates the 5% Bonferroni threshold. (D) Difference in mean raw reflectance between the high- and low-boron groups (high minus low). Shaded bands indicate ±1 SE of the difference. (E) Wavelength-wise Pearson correlations between reflectance measured before treatment and at days of treatment (DOT) 5, 13, and 21, and boron BLUEs derived from the F_1_ population (n = 162). Panels A and D show raw reflectance spectra, whereas panels B, C, and E show analyses based on standard normal variate (SNV)-transformed reflectance.

In the F_1_ population, correlations between reflectance measured at each time point and leaf boron concentration at harvest changed during the experiment (Figure 5E). Before treatment, reflectance near 682 nm was positively correlated with leaf boron concentration (r = 0.39), whereas reflectance near 563-584 nm was negatively correlated (r = −0.39). During boron treatment, the strongest positive correlation increased to r = 0.62 on DOT 5 and 13 before decreasing to r = 0.54 on DOT 21. The strongest negative correlation followed a similar pattern, increasing in magnitude to r = −0.62 during treatment before decreasing to r = −0.50 on DOT 21. Correlations across the near-infrared region were close to zero on DOT 21 (mean r = −0.03 across 780-880 nm).

## Discussion

Leaf boron accumulation varied approximately 190-fold among accessions in the diversity panel, and entry-mean heritability reached H² = 0.71, indicating a strong genetic component. Broad genetic variation in ion accumulation has also been reported for chloride and sodium in *Vitis* (Heinitz et al., 2020; Henderson et al., 2018; Sharma et al., 2025), and for boron in several species (Jia et al., 2021; Nable et al., 1997; Tergemina et al., 2026). This variation occurred both between and within species, with *V. acerifolia* and *V. riparia* having lower concentrations than *V. arizonica*, *V. candicans*, and *V. champinii* (Figure 1D). The substantial within-species variation observed for all three traits suggests that useful low-boron alleles may segregate within individual species. This is important for breeding boron-tolerant rootstocks because low-boron haplotypes may be recoverable within rootstock-relevant germplasm, such as *V. rupestris* and *V. riparia*, which have historically, and in recent studies, shown superior rooting ability (Heinitz et al., 2019; Sharma et al., 2026), without requiring wide interspecific hybridization. A comparable pattern has been reported for chloride exclusion, where substantial variation occurs among accessions within species (Heinitz et al., 2020; Sharma et al., 2025).

Stomatal conductance was positively correlated with leaf boron concentration (r = 0.34, P = 1.31 × 10⁻⁸; Figure 1G), consistent with the expectation that higher transpiration contributes to greater delivery of xylem-mobile boron to leaves (Brown & Shelp, 1997; Miwa & Fujiwara, 2010). This moderate correlation, with g_s_ explaining approximately 12% of the variance in leaf boron (r² = 0.12), indicates that transpiration-driven delivery contributes to leaf boron accumulation but explains only part of the genotypic variation. This interpretation is consistent with observations in barley, where differences in water-use efficiency between boron-tolerant and boron-sensitive genotypes were insufficient to explain the much larger differences in boron accumulation (Nable et al., 1997). Other processes, including root uptake, xylem loading, cell wall retention, and internal partitioning, are likely to contribute additional genotypic variation (Miwa & Fujiwara, 2010; O’Neill et al., 2004). Leaf boron concentration was weakly but positively correlated with Φ_PSII_ (r = 0.20, P = 8.16 × 10⁻⁴; Figure 1H), so higher boron was not associated with reduced Φ_PSII_ across the diversity panel. This pattern is more consistent with Lupo et al. (2025), who found no overall relationship between leaf boron concentration and Φ_PSII_, than with Pech et al. (2013), who reported reduced Φ_PSII_ under excess boron. Differences among studies may reflect variation in supplied boron concentration, exposure duration, genotypes examined, or the timing of physiological measurements relative to stress development. Nevertheless, excess boron can induce oxidative stress in grapevine (Gunes et al., 2006) and other species (Cervilla et al., 2007), indicating that physiological injury may occur without a consistent decline in Φ_PSII_.

To account for variation in transpiration-related boron delivery (Brown & Shelp, 1997; Miwa & Fujiwara, 2010), gₛ was included as a covariate in the boron GWAS. The chromosome 11 association at 2.33 Mb was detected at the same position without gₛ adjustment (P = 3.23 × 10⁻⁷; data not shown) and showed stronger statistical support after adjustment (P = 1.12 × 10⁻⁷; Figure 2A), indicating that the signal was not driven solely by variation in gₛ. The nearest significant gₛ association was approximately 190 kb away, at 2.14 Mb. In a previous analysis of the same *Vitis* germplasm collection, linkage disequilibrium decayed rapidly, reaching half its maximum average r² within 2.12 kb (Cochetel et al., 2023), substantially shorter than the distance between these associations. The two signals are therefore likely to represent distinct loci despite their physical proximity.

The suggestive chromosome 11 GWAS signal pointed to a candidate region associated with leaf boron accumulation after accounting for g_s_. Although the GWAS association was only suggestive, QTL mapping in the F_1_ population independently identified a significant chromosome 11 QTL (LOD = 7.34) that explained 19.5% of the phenotypic variance. Projection between the two reference assemblies placed the GWAS and QTL peaks within the same syntenic region despite a 200-330-kb difference between their projected positions. Given the proportion of phenotypic variance explained and its potential relevance for breeding, we designate this QTL as qBEx11.1. The convergence of the two mapping approaches prioritizes this region for mapping, validation, and deployment.

At the qBEx11.1 peak, the Longii-9018 B haplotype and GRN3 D haplotype were associated with higher leaf boron accumulation, whereas the AC genotype class, combining the lower-boron haplotypes from both parents, had the lowest mean (Figure 3C, D). This approximately additive pattern suggests that markers linked to qBEx11.1 may be useful for selecting lower-boron progeny within related families, following validation in additional populations and environments.

The ±50-kb interval surrounding the suggestive GWAS peak at chr11:2.33 Mb contained two preliminary positional candidates: a probable pectinesterase 53 approximately 13 kb from the lead SNP and a MATE efflux family protein approximately 33 kb away (Table S2). The two closest annotated genes, encoding a COMPASS-like WDR5A component approximately 0.5 kb away and a probable glutamate carboxypeptidase approximately 1.3 kb away, are not known to have direct roles in boron homeostasis. We therefore prioritized the pectinesterase and MATE genes based on their functional plausibility.

The pectinesterase represents a biologically plausible candidate because boron contributes to plant cell wall structure through borate diester cross-linking of RG-II, a pectic polysaccharide required for normal wall structure and growth (Goldbach & Wimmer, 2007; O’Neill et al., 2004). Pectin methylesterases modify the methylesterification of pectin and thereby influence its charge and the physical properties of the cell wall (Pelloux et al., 2007). Although a direct role for pectinesterases in boron homeostasis has not been established, variation in pectin modification could influence leaf boron retention through changes in cell wall organization.

NIP boric-acid channels and BOR borate exporters have established roles in boron uptake and distribution in plants (Sabir et al., 2020; Takano et al., 2002, 2006; Tanaka et al., 2008). No NIP or BOR genes were annotated within the ±50-kb interval surrounding the chromosome 11 lead SNP. The interval contained a gene annotated as a MATE efflux family protein. Plant MATE proteins transport diverse substrates, including organic acids involved in mineral-stress responses and detoxification (Magalhães et al., 2007; Yokosho et al., 2016), and three clustered *Arabidopsis* MATE genes were strongly induced in leaves under boron toxicity (Kasajima & Fujiwara, 2007). These findings make the MATE gene a plausible positional candidate but do not establish a direct role in boron transport. Fine mapping, expression analysis, and functional validation will be required to determine whether the pectinesterase, MATE transporter, or another gene underlies qBEx11.1.

The two strongest gₛ loci contained positional candidates for stomatal regulation. At chr18:5.94 Mb, the lead SNP occurred within a gene encoding a GATA transcription factor, and a second GATA-type zinc-finger gene was located approximately 17 kb away (Table S2). Plant GATA transcription factors regulate several light-responsive and developmental processes, including chloroplast development and, for some B-GATA family members, stomatal development (Behringer & Schwechheimer, 2015; Klermund et al., 2016). At chr3:2.55 Mb, the lead SNP was located approximately 1.6 kb from an L-ascorbate oxidase homolog. Ascorbate oxidase influences the redox state of the apoplast and has been associated experimentally with changes in g_s_ (Pignocchi et al., 2003; Fotopoulos et al., 2008). These annotations identify candidates for further investigation but do not establish their causal roles in variation in gₛ.

Boron toxicity and salinity frequently co-occur in irrigated arid soils, but their interaction varies among species and conditions. Salinity reduced leaf boron accumulation in grapevine (Yermiyahu et al., 2007), whereas both mitigation and intensification of boron toxicity have been reported in other systems (Pandey et al., 2019). This co-occurrence raises the practical question of whether genotypes selected for low boron accumulation also accumulate less chloride. Leaf boron and chloride were moderately correlated (r = 0.36, P = 1.03 × 10⁻⁸; Figure 4A). Although 116 of the 118 SNPs suggestive for either trait had effects in the same direction (Figure 4C), none was suggestive for both traits, and no 500-kb window contained suggestive associations for both (Figure 4B, D). Thus, the matched GWAS analyses provided no evidence of a shared major locus. This interpretation is supported by the chloride QTL results from the same Longii-9018 × GRN3 F_1_ population. The chloride exclusion QTL qClEx8.1 mapped to chromosome 8 (Sharma et al., 2025), rather than chromosome 11, where qBEx11.1 was identified. Moreover, because both traits were mapped in the same population, the absence of overlap between the boron and chloride QTLs is not attributable to differences in mapping population. Together, these findings support largely distinct genetic control of boron and chloride accumulation. Selecting for boron exclusion may not reliably reduce leaf chloride accumulation, or vice versa. Genotypes with low accumulation of both boron and chloride are promising candidates for further validation and may be useful as crossing parents, provided that these traits are stable across environments and compatible with other rootstock characteristics. We therefore suggest that both ions be measured directly when evaluating rootstocks for environments affected by both boron and salinity.

Leaf spectral reflectance was associated with leaf boron in both the wild *Vitis* diversity panel and the F_1_ population. Significant wavelength intervals were concentrated in the visible and red-edge regions in both populations and extended into the near-infrared in the F_1_ population (Figure 5B, C). These intervals include spectral bands sensitive to chlorophyll content and red-edge position (Horler et al., 1983; Sims & Gamon, 2002), suggesting that the observed associations may reflect variation in pigment status or other leaf properties related to boron accumulation. Spectral correlations with leaf boron were detectable before treatment, became strongest at DOT 5 and 13, and weakened by DOT 21. The most affected leaves showed advanced toxicity symptoms at harvest, including chlorosis, necrosis, and marginal scorch (Figure S1), consistent with reported symptoms of boron toxicity (Nable et al., 1997; Yermiyahu et al., 2007). Tissue injury may have had a greater influence on leaf reflectance late in the experiment, although symptom severity was not quantified. The stronger correlations at DOT 5 and 13 suggest that measurements made before severe injury develops may provide a more informative phenotyping window. The present analysis evaluated spectral associations rather than predictive performance. Cross-validated models are therefore needed to determine whether compact leaf spectrometry can support germplasm screening before destructive elemental analysis.

## Summary

In summary, the high heritability of leaf boron accumulation and the convergence of QTL and association mapping on qBEx11.1 show that natural variation in boron accumulation is genetically tractable in wild *Vitis*. The modest relationship with gₛ and the distinct genomic associations for chloride suggest that this variation is not simply a consequence of transpiration or of a general capacity to restrict ion uptake. These findings establish qBEx11.1 as a priority for validation across populations and environments. The boron-associated reflectance signals provide a complementary approach for more efficient phenotyping, although cross-validated prediction models are needed before they can be applied in germplasm screening. Together, genetic markers and spectral phenotyping could improve the selection of boron-tolerant grapevine rootstocks.

## Acknowledgments

The authors would like to thank Paul Skinner for the initial discussions and conceptualization of this study. This project was partially funded by the American Vineyard Foundation (project 2025-2953), the California Grape Rootstock Improvement Commission, the California Grape Rootstock Research Foundation, the USDA National Institute of Food and Agriculture Specialty Crop Research Initiative (2024-51181-43236 and 2026-67013-45985), and the National Science Foundation grant #1741627. This project was supported by BARD, the United States–Israel Binational Agricultural Research and Development Fund, Vaadia-BARD Postdoctoral Fellowship Award No FI-633-2023.

## Supplemental Material

The supplemental materials include two figures showing representative boron toxicity symptoms and QTL mapping results for the Longii-9018 × GRN3 F_1_ population. Two tables summarize the datasets and sample sizes used and list annotated genes near selected leaf-boron and stomatal-conductance lead SNPs.

## Data Availability

Raw sequencing reads are available in the NCBI Sequence Read Archive under BioProject accessions PRJNA731597, PRJNA842753, and PRJNA984685. The chloride phenotype data used in the comparative analyses are available in Supplementary Table 2 of Sharma et al. (2025).

## Author Contributions

**YL**: Conceptualization; Data Curation; Formal Analysis; Investigation; Methodology; Project Administration; Software; Validation; Visualization; Writing-original draft; Writing-review and editing. **SS**: Investigation; Methodology. **JRM**: Investigation. **VN**: Investigation; Methodology; Resources. **AG**: Investigation; Methodology; Resources. **NC**: Data Curation; Formal Analysis; Resources; Writing-review and editing. **DC**: Data Curation; Formal analysis; Resources; Writing-review and editing. **AJM**: Conceptualization; Writing-review and editing. **LDG**: Conceptualization; Data Curation; Formal Analysis; Funding Acquisition; Investigation; Methodology; Project Administration; Software; Supervision; Validation; Visualization; Writing-review and editing.

## Abbreviations

BH: Benjamini-Hochberg
BLUE: best linear unbiased estimate
chr: chromosome
DOT: day of treatment
DW: dry weight
F_1_: first filial generation
FDR: false discovery rate
GATK: Genome Analysis Toolkit
GEMMA: genome-wide efficient mixed-model association
g_s_: stomatal conductance
GWAS: genome-wide association study
LOD: logarithm of odds
MAF: minor allele frequency
PSII: photosystem II
Φ_PSII_: photosystem II quantum yield
QTL: quantitative trait locus
REML: restricted maximum likelihood
SNP: single-nucleotide polymorphism
SNV: standard normal variate.

