## supplemental materials for "Genetic Mapping Reveals a Major Locus for Boron Exclusion in Wild *Vitis*"

**Number of pages:** 6

**Number of supplemental figures:** 2

**Number of supplemental tables:** 2

**Contents**

**Supplemental Figure S1.** Boron toxicity symptoms in the Longii-9018 × GRN3 $F_1$ mapping population at harvest.

**Supplemental Figure S2.** Parent-specific QTL scans and linkage-map summary for leaf boron concentration in the Longii-9018 × GRN3 $F_1$ population.

**Supplemental Table S1.** Sample sizes and datasets used for phenotypic, GWAS, QTL, and spectral analyses.

**Supplemental Table S2.** Annotated genes within ±50 kb of the chromosome 11 lead SNP for leaf boron concentration and three selected lead SNPs for stomatal conductance.

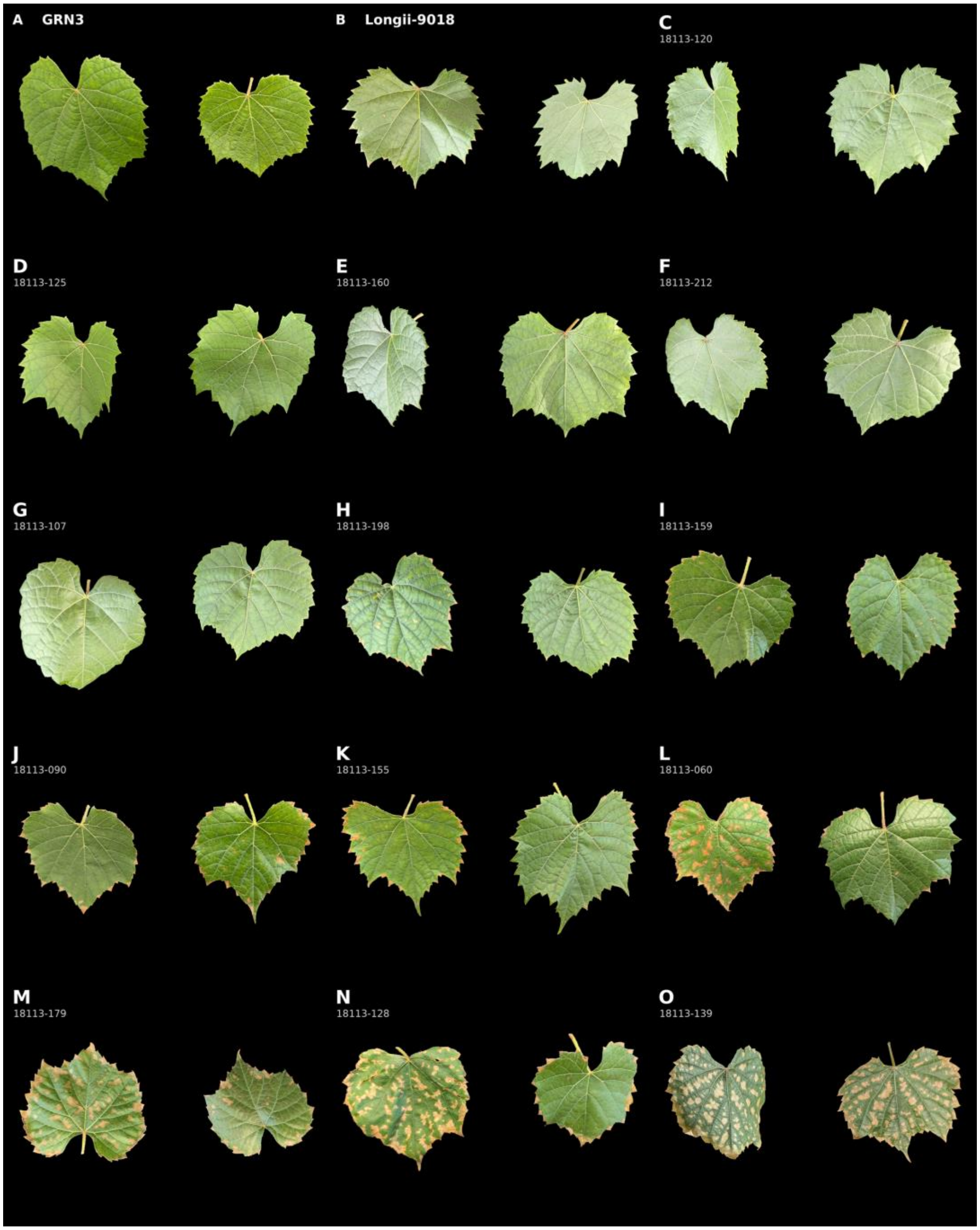


**Supplemental Figure S1. Boron toxicity symptoms in the Longii-9018 × GRN3 $F_1$ mapping population at harvest.** Representative leaves photographed after boron treatment. **(A)** GRN3 parent; **(B)** Longii-9018 parent; **(C-O)** $F_1$ progeny arranged by increasing symptom severity. Symptoms ranged from interveinal chlorosis and marginal scorch to necrotic lesions and differed among progeny, illustrating variation in the boron-toxicity response. Each panel shows two leaves from the same genotype.

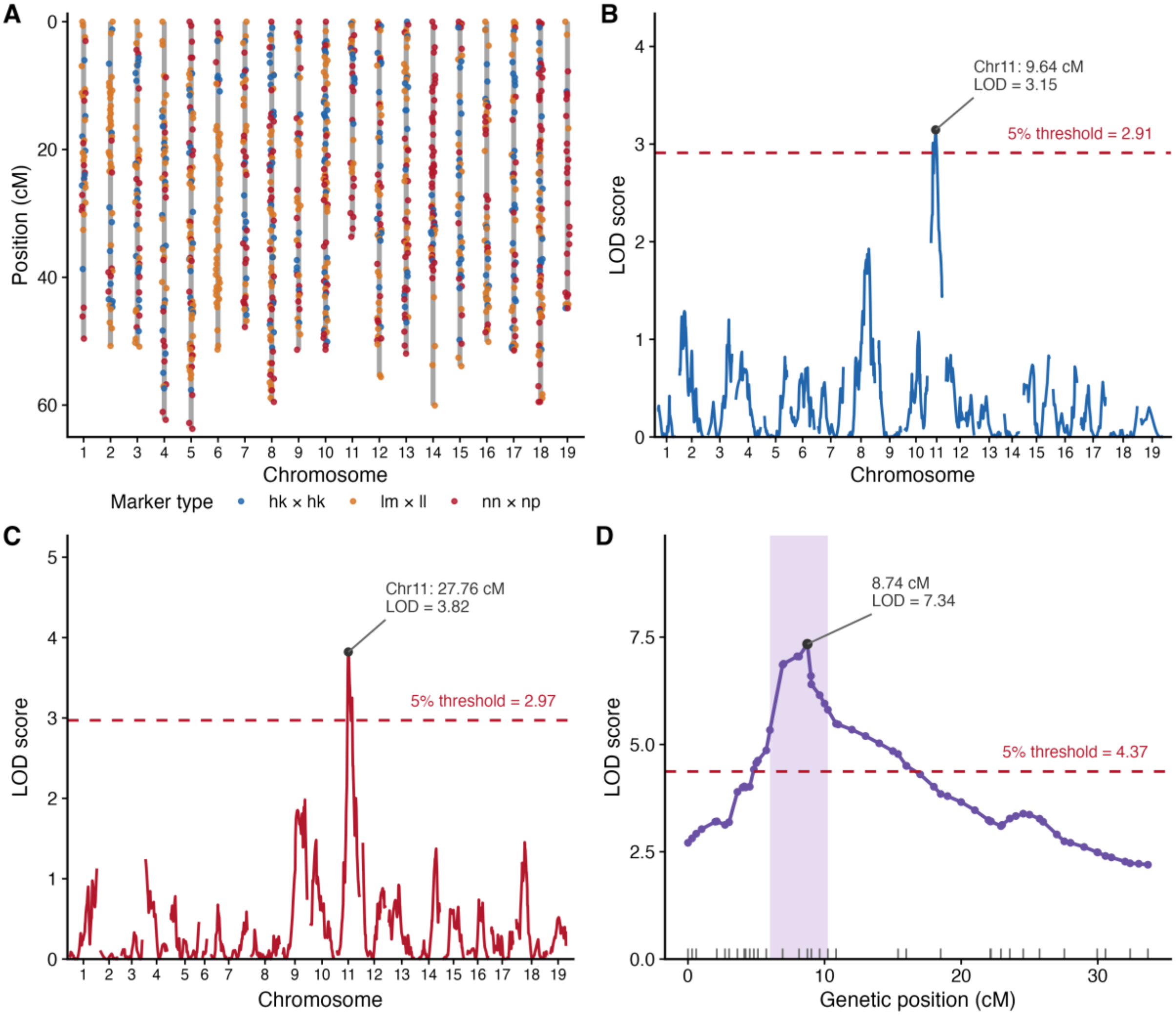


**Supplemental Figure S2. Parent-specific QTL scans and linkage-map summary for leaf boron concentration in the Longii-9018 × GRN3 $F_1$ population.** (A) Composite genetic map containing 1038 markers across 19 chromosomes; points indicate marker positions and segregation classes. Parent-specific scans of leaf boron BLUEs were performed using (B) the Longii-9018 map and (C) the GRN3 map. Each parent-specific scan was analyzed as a backcross. The maximum LOD scores occurred on chromosome 11 at 9.64 cM for Longii-9018 (LOD = 3.15; threshold = 2.91) and 27.76 cM for GRN3 (LOD = 3.82; threshold = 2.97). Dashed red lines indicate the 5% genome-wide significance thresholds determined separately for each scan using 1000 permutations. Filled points, leader lines, and labels identify the maximum LOD positions. (D) Chromosome 11 LOD profile from the primary four-way scan of leaf boron BLUEs. Purple points and the connecting line show the LOD profile, and the dark point identifies the maximum LOD score at 8.74 cM (LOD = 7.34). The shaded region indicates the 1.5-LOD support interval from 6.0 to 10.2 cM, the dashed red line indicates the 5% genome-wide permutation threshold (LOD = 4.37), and short vertical lines along the x-axis indicate observed marker positions.

**Supplemental Table S1.** Sample sizes and datasets used for phenotypic, genome-wide association, quantitative trait locus, and spectral analyses. Sample sizes differ because each analysis required a different combination of phenotypic, single-nucleotide polymorphism, covariate, marker, or reflectance data.

| **Analysis** | **n** | **Dataset and inclusion criterion** |
|---|---|---|
| ***Vitis* diversity panel** | | |
| Leaf boron BLUEs and heritability | 267 | Heritability estimated from 775 observations (mean 2.9 per accession). |
| $g_s$ BLUEs and heritability | 265 | Heritability estimated from 973 observations (mean 3.7 per accession). |
| $\Phi_{PSII}$ BLUEs | 265 | Accessions with usable physiological measurements. |
| Leaf boron concentration GWAS (main) | 224 | Boron, $g_s$, and SNP data; $g_s$ included as a covariate. |
| Leaf boron concentration GWAS (sensitivity analyses) | 224 | Same accessions and SNPs as the main GWAS; $g_s$ omitted. |
| $g_s$ GWAS | 233 | Accessions with $g_s$ and SNP data. |
| Leaf boron-chloride phenotypic comparison | 241 | Accessions with measurements for both traits. |
| Matched boron-chloride GWAS | 222 | The same accessions and 1,703,430 SNPs were used for both traits. |
| Spectral analysis | 238 | Accessions with matched leaf boron and reflectance data. |
| **Longii-9018 × GRN3 $F_1$ population** | | |
| Leaf boron concentration distribution | 162 | Progeny with leaf boron BLUEs; parental BLUEs shown as references. |
| Leaf boron concentration heritability | 162 | Estimated from 486 observations (three per progeny); parents excluded. |
| Leaf boron concentration QTL mapping | 158 | Progeny with leaf boron BLUEs and marker data; used in the four-way and parent-specific scans. |
| Highest-LOD marker genotype comparison | 156 | Progeny with nonmissing genotypes at the chromosome 11 marker. |
| Spectral analysis | 162 | Progeny with matched leaf boron and reflectance data. |
| **GWAS genotype-class comparisons** | | |
| Leaf boron concentration, Chr11:2.33 Mb (Fig. 2D) | 212 | Non-missing genotypes at the lead SNP: TT, 104; CT, 58; CC, 50. |
| $g_s$, Chr18:5.94 Mb (Fig. 2H) | 209 | Non-missing genotypes at the lead SNP: TT, 88; CT, 61; CC, 60. |
| $g_s$, Chr3:2.55 Mb (Fig. 2I) | 201 | Non-missing genotypes at the lead SNP: TT, 156; GT, 36; GG, 9. |
| $g_s$, Chr11:2.14 Mb (Fig. 2J) | 209 | Non-missing genotypes at the lead SNP: GG, 175; AG, 21; AA, 13. |

*Abbreviations:* BLUE, best linear unbiased estimate; GWAS, genome-wide association study; QTL, quantitative trait locus; SNP, single-nucleotide polymorphism; $g_s$, stomatal conductance; $\Phi_{PSII}$, photosystem II quantum yield.

**Supplemental Table S2.** Annotated genes within ±50 kb of the chromosome 11 lead single-nucleotide polymorphism for leaf boron concentration and three selected lead single-nucleotide polymorphisms for stomatal conductance.

| Gene ID | Coordinates | Strand | Distance | Annotation |
|---|---|---|---|---|
| — | bp | — | kb | — |
| **Leaf boron concentration: chr11:2.33 Mb (region also supported by the $F_1$ QTL)** | | | | |
| chr11.ver1.0.g183300 | 2,279,939-2,283,738 | + | −43.8 | mitogen-activated protein kinase kinase 2-like |
| chr11.ver1.0.g183310 | 2,285,822-2,287,082 | + | −40.5 | oxidative stress 3 |
| chr11.ver1.0.g183320 | 2,294,101-2,308,130 | + | −19.4 | Arginyl-tRNA synthetase, class Ic |
| chr11.ver1.0.g183330 | 2,309,101-2,316,680 | + | −10.9 | phosphoinositide 3-kinase regulatory subunit 4-like |
| chr11.ver1.0.g183340 | 2,316,755-2,327,042 | + | −0.5 | COMPASS-like H3K4 histone methylase component WDR5A |
| chr11.ver1.0.g183350 | 2,328,845-2,336,576 | - | +1.3 | probable glutamate carboxypeptidase 2 |
| **chr11.ver1.0.g183360**[a] | **2,340,431-2,343,563** | **+** | **+12.9** | **probable pectinesterase 53** |
| chr11.ver1.0.g183370 | 2,344,947-2,348,202 | - | +17.4 | protein IQ-DOMAIN 14-like |
| chr11.ver1.0.g183380 | 2,352,470-2,355,690 | + | +24.9 | nuclear poly(A) polymerase 4-like isoform X4 |
| **chr11.ver1.0.g183390** | **2,360,667-2,362,416** | **-** | **+33.1** | **MATE efflux family protein** |
| chr11.ver1.0.g183400 | 2,370,795-2,372,739 | + | +43.2 | hypothetical protein L484_006995 |
| chr11.ver1.0.g183410 | 2,376,076-2,383,827 | + | +48.5 | hexokinase |
| **Stomatal conductance ($g_s$): chr18:5.94 Mb** | | | | |
| chr18.ver1.0.g285570 | 5,917,342-5,918,952 | - | −36.4 | protein Rf1, mitochondrial-like isoform X1 |
| **chr18.ver1.0.g285580** | **5,922,055-5,938,718** | **+** | **−16.6** | **GATA-type zinc finger protein with TIFY domain-containing protein** |
| **chr18.ver1.0.g285590** | **5,940,577-5,962,177** | **+** | **0.0** | **GATA transcription factor** |
| chr18.ver1.0.g285600 | 5,962,757-6,058,816 | - | +7.4 | protein SABRE-like isoform X1 |
| chr18.ver1.0.g285610 | 5,966,728-5,970,980 | - | +11.4 | hypothetical protein PRUPE_ppa021750mg, partial |
| **Stomatal conductance ($g_s$): chr3:2.55 Mb** | | | | |
| chr03.ver1.0.g060090 | 2,501,532-2,506,796 | + | −43.7 | SAUR-like auxin-responsive protein family |
| chr03.ver1.0.g060100 | 2,509,021-2,509,629 | + | −40.8 | DUF4283 domain protein |
| chr03.ver1.0.g060110 | 2,509,702-2,511,443 | + | −39.0 | endonuclease/exonuclease/phosphatase family protein |
| chr03.ver1.0.g060120 | 2,513,288-2,514,112 | + | −36.3 | homogentisate geranylgeranyltransferase-like |
| chr03.ver1.0.g060130 | 2,516,390-2,520,050 | + | −30.4 | lysine histidine transporter-like 8 |
| chr03.ver1.0.g060140 | 2,522,879-2,523,920 | - | −26.5 | Tetraacyldisaccharide 4'-kinase |
| chr03.ver1.0.g060150 | 2,535,807-2,537,928 | + | −12.5 | lysine histidine transporter-like 8 |
| chr03.ver1.0.g060160 | 2,539,385-2,540,032 | + | −10.4 | dehydrin family protein |
| chr03.ver1.0.g060170 | 2,541,076-2,543,927 | - | −6.5 | late embryogenesis abundant protein |
| chr03.ver1.0.g060180 | 2,546,069-2,547,623 | - | −2.8 | ankyrin repeat-containing protein 2 |
| **chr03.ver1.0.g060190** | **2,552,052-2,555,444** | **+** | **+1.6** | **L-ascorbate oxidase homolog** |
| chr03.ver1.0.g060200 | 2,559,090-2,561,255 | + | +8.6 | GDSL esterase/lipase At5g55050-like |
| chr03.ver1.0.g060210 | 2,560,395-2,564,209 | - | +9.9 | adenine nucleotide alpha hydrolase-like domain kinase |

| Gene ID | Coordinates | Strand | Distance | Annotation |
|---|---|---|---|---|
| — | bp | — | kb | — |
| chr03.ver1.0.g060220 | 2,565,048-2,570,270 | - | +14.6 | coiled-coil domain-containing protein 124 |
| chr03.ver1.0.g060230 | 2,574,380-2,582,606 | + | +23.9 | Rop guanine nucleotide exchange factor 1 |
| chr03.ver1.0.g060240 | 2,577,566-2,582,341 | - | +27.1 | F-box SKIP23-like protein |
| chr03.ver1.0.g060250 | 2,583,765-2,587,302 | + | +33.3 | phosphoglycerate mutase |
| chr03.ver1.0.g060260 | 2,588,610-2,590,902 | + | +38.2 | small nuclear ribonucleoprotein SmD3b-like |
| chr03.ver1.0.g060270 | 2,590,954-2,593,705 | - | +40.5 | armadillo/beta-catenin repeat family protein |
| **Stomatal conductance ($g_s$): chr11:2.14 Mb (approximately 190 kb from the boron lead SNP)** | | | | |
| chr11.ver1.0.g183090 | 2,091,240-2,095,165 | - | −47.4 | IAA-amino acid hydrolase ILR1-like protein |
| chr11.ver1.0.g183100 | 2,100,829-2,105,626 | + | −36.9 | vacuolar protein sorting-associated protein 32 homolog 2-like |
| chr11.ver1.0.g183110 | 2,111,484-2,112,096 | - | −30.5 | arabinogalactan peptide 22-like |
| chr11.ver1.0.g183120 | 2,113,207-2,123,711 | - | −18.9 | Mnd1 family protein |
| chr11.ver1.0.g183130 | 2,129,149-2,130,696 | - | −11.9 | transcription factor RAX2-like |
| chr11.ver1.0.g183140 | 2,134,914-2,137,574 | - | −5.0 | zinc finger CCCH domain-containing protein 2-like |
| chr11.ver1.0.g183150 | 2,152,935-2,155,260 | + | +10.4 | myo-inositol oxygenase |
| chr11.ver1.0.g183160 | 2,158,346-2,187,680 | + | +15.8 | glycine-rich protein |
| chr11.ver1.0.g183170 | 2,189,897-2,193,942 | + | +47.3 | gamma-glutamyltranspeptidase 1-like |

*Note:* Coordinates and annotations are from haplotype 1 of the *Vitis girdiana* SC2 reference genome (VITVgdSC2_v1.0). Distance is measured from the lead SNP to the nearest gene boundary. Negative values indicate genes located at lower genomic coordinates than the lead SNP, positive values indicate genes at higher genomic coordinates, and zero indicates that the lead SNP lies within the gene. Gene IDs omit the common prefix VITVgdSC2_v1.0.hap1. Candidate assignments are positional and require fine mapping, expression analysis, and functional validation.

*Abbreviations:* $F_1$, first filial generation; GWAS, genome-wide association study; QTL, quantitative trait locus; SNP, single-nucleotide polymorphism; $g_s$, stomatal conductance.

[a] Boldface identifies positional candidates discussed in the manuscript.
